# Maturation of Cognitive Control in the Inferior Frontal Junction: A Combined Systematic Review and Coordinate-Based fMRI Meta-Analysis

**DOI:** 10.64898/2026.05.01.722218

**Authors:** Cidney Montford, Jeremy Hogeveen

## Abstract

Cognitive control is fundamental to goal-directed behavior, and its protracted maturation is a hallmark of adolescent brain development. In adulthood, the inferior frontal junction (IFJ) is functionally characterized as a critical region for updating task representations to guide the implementation of cognitive control. Yet, how its domain-general control functions emerge and mature across development remains largely underexplored. Specifically, it is unclear whether the IFJ’s capacity for cognitive control enhances uniformly as a single construct, or if this region matures asynchronously for distinct control processes like inhibition, switching, and working memory. To address this gap, we conducted a combined systematic review and coordinate-based neuroimaging meta-analysis. Applying multilevel kernel density analyses to fMRI studies of inhibition, switching, and working memory in youth and adults, we synthesized data from 72 contrasts (779 foci; N = 1,913). The results revealed a staggered developmental trajectory for IFJ recruitment. While adults showed consistent convergence of activation in the IFJ across all three domains, youth exhibited robust bilateral IFJ convergence exclusively during inhibitory control tasks. This suggests inhibition may be a developmentally foundational process localized to this region earlier in the lifespan. Furthermore, adults demonstrated hemispheric specialization absent in youth: left IFJ was uniquely sensitive to switching and working memory in adults, but not in youth. Together, these findings support a model where the IFJ does not mature as a static, monolithic node, but rather acts as a dynamic hub that integrates fractionated cognitive processes at different stages of development.

## Introduction

The ability to flexibly adapt our thoughts and behaviors to changing goals—broadly termed ‘cognitive control’—is a defining and remarkable feature of the human brain. While the protracted development of cognitive control from childhood through adulthood is well-documented behaviorally, the exact neurocognitive mechanisms driving this maturation are a critical open question for the field. Here, we focus on a specialized node of the human prefrontal cortex—the inferior frontal junction (IFJ). Situated at the anatomical intersection of attention, motor, and working memory pathways, the IFJ is thought to serve as a “functional hub” for the frontoparietal network, flexibly updating task representations and coordinating multiple domains of cognitive control (Brass et al., 2005; Derrfuss et al., 2005; Dixon et al., 2018). Yet, a fundamental question regarding its ontogeny remains: How and when does IFJ’s domain-general functional role emerge across neurodevelopment? It is currently unclear whether the IFJ develops as a monolithic node, or whether its functional architecture develops to subserve specific cognitive control functions at different stages of the lifespan.

To resolve this, the current review combines a comprehensive synthesis of the literature and a coordinate-based neuroimaging meta-analysis to map IFJ recruitment across three core domains of cognitive control—inhibitory control, switching, and working memory—in both youth and adults. Our approach traces the trajectory of IFJ’s functional maturation to provide a novel framework for understanding the extent to which cognitive control operations might rely on common or distinct functional networks anchored in the IFJ across development. Before we describe the current fMRI meta-analysis and its implications, we first operationalize the components of cognitive control and review the distinct functional role that IFJ appears to play within the mature human brain.

### The three-factor model of cognitive control

“Executive functions” (EFs) are a general set of cognitive functions involved in the planning and regulation of goal-directed behaviors, which tend to be disrupted in patients with prefrontal cortex damage (Cristofori et al., 2019). While profoundly useful from clinical and neurodevelopmental perspectives, EFs have been difficult to pin down mechanistically using cognitive computational modeling and/or neuroimaging approaches. This difficulty has led much of the field to shift towards the preferred term “cognitive control,” referring to three component processes or subdomains: inhibition, task-switching or set-shifting (i.e., cognitive flexibility), and working memory (McKenna et al., 2017; Miyake et al., 2000; Sundermann & Pfleiderer, 2012).

Miyake et al.’s (2000) classic three-factor model of EFs denotes both unity and diversity of the inhibition, flexibility, and working memory components of cognitive control. While their confirmatory factor analysis indicates each domain measures different components of cognitive control (diversity), a further test of independence suggests some commonality, or unity, among the functions—an overlapping component they referred to as the “common executive” (McKenna et al., 2017; Miyake et al., 2000). The comprehensive three-factor model proposed by Miyake and colleagues (2000) is widely used, and has been replicated in both adults (Karr et al., 2018) and youth (McKenna et al., 2017). While McKenna and colleagues (2017) report a three-factor structure in youth above the age of six, a recent re-analysis by Karr et al. (2018) found a two-factor model yielded a better fit in youth. Specifically, shifting merged with working memory and shifting merged with inhibition both produced better fits than the three-factor model in children and adolescents (Karr et al., 2018). This discrepancy suggests that the flexibility component of cognitive control may mature more slowly than inhibition or working memory, emerging as a distinct domain of control later in development.

To operationalize cognitive control in a way that is relevant for the current fMRI meta-analysis, we focus on the specific tasks most often used to isolate subdomains of cognitive control in the scanner. Inhibitory control involves the suppression of automatic or prepotent responses (Miyake et al., 2000), and is frequently measured in fMRI using anti-saccade, flanker, go/no-go, stop signal, and Stroop tasks (Z. Zhang et al., 2021). Despite known construct validity nuances regarding inhibitory demands in standard go/no-go paradigms (Wessel, 2018), we include these tasks because they are the modal approach used in developmental neuroimaging studies, maximizing the statistical power of the current meta-analysis. Task-switching or set-shifting paradigms are general approaches to operationalizing cognitive flexibility in the scanner—comprising the ability to engage and disengage task sets or shift mental sets in spite of proactive interference (Kim et al., 2011; Miyake et al., 2000). This domain is typically measured using tasks that involve alternating rule- or context-dependent shifts in stimulus-response mappings. Finally, working memory—often used interchangeably with “updating” in cognitive control and EF models—entails the active maintenance of task-relevant information in line with one’s goals (Miyake et al., 2000), and is most often operationalized using variants of the *N*-back paradigm (Z. Zhang et al., 2021).

### The inferior frontal junction

The IFJ is a key component of the frontoparietal network (FPN), a neural circuit which is heavily involved in cognitive control (Dixon et al., 2018; Keller et al., 2023). The Glasser atlas parcellation defines the region with two sub-regions: the anterior IFJ (IFJa) and the posterior IFJ (IFJp; See **Figure 1**; Glasser et al., 2016). The IFJ is of special interest given its functional network architecture. Using the Glasser atlas parcellation to examine functional connectivity patterns, the IFJ region exhibits hemispheric differences, whereby the right IFJp is shown to be a part of the FPN, while the left IFJp is shown to be at the intersection of the FPN and the DAN; bilaterally, the IFJa is shown to be a part of the FPN (Glasser et al., 2016; Schaefer et al., 2018). Depending on the specific tasks and analysis approaches, it has been found to be functionally connected to the cingulo-opercular, default, frontoparietal, ventral attention/salience, and dorsal attention networks in a flexible, task-dependent manner (Asplund et al., 2010; Corradi-Dell’Acqua et al., 2015; Dixon et al., 2018; Roth et al., 2014; Yin et al., 2018). The functional network flexibility of the IFJ is due to its location at the junction of the inferior frontal sulcus (IFS) and the inferior precentral sulcus (IPCS; Brass et al., 2005; Derrfuss et al., 2005). By bridging the IFS—which is implicated in abstract encoding of goal-relevant information in working memory (Nolan et al., 2024; Ruland et al., 2022)—and the IPCS—which borders motor control regions and is also involved in visual selective attention as it contains the “premotor eye fields” (PEF; Glasser et al., 2016; Noyce et al., 2017)—the IFJ serves as a convergence zone for multiple functionally heterogeneous neuronal circuits relevant for visual attention and cognitive control (Brass et al., 2005). Crucially, the IFJ’s unique position between brain regions involved in higher-order abstract processing and concrete sensorimotor planning and action selection makes the IFJ ideally placed to mediate the core domains of cognitive control. The IFJ translates internal goals into active task sets (i.e., supporting working memory), dynamically updates these representations when task-relevant rules or contexts change (i.e., supporting cognitive flexibility), and gates downstream motor output to enable suppression of inappropriate prepotent actions (i.e., supporting inhibitory control; Levy & Wagner, 2011). Effective connectivity analyses have found IFJ to be predictive of activity in other regions during cognitive control tasks, leading to the suggestion that it acts as a “switchboard” or hub between functional networks (Hippmann et al., 2021; Sneve et al., 2013; Yin et al., 2018).

**Figure 1.**
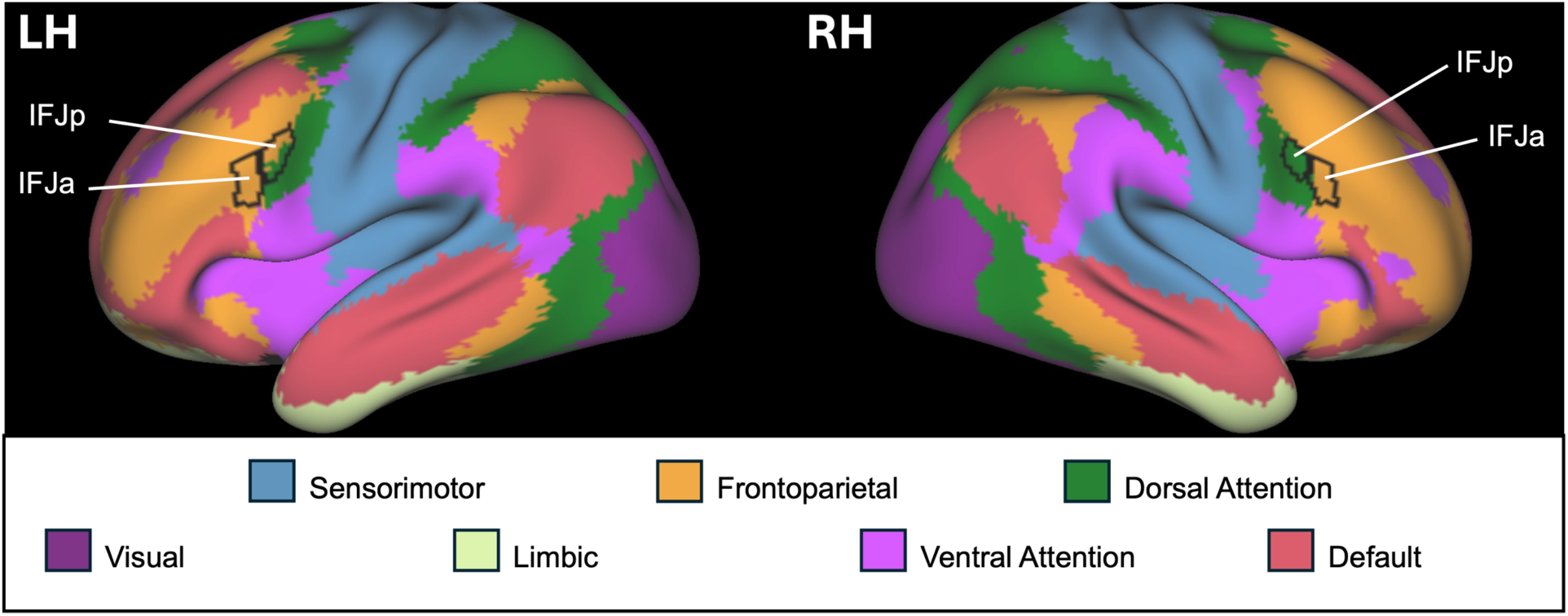
Inferior Frontal Junction location from Glasser (2016) atlas, and functional networks from Schafer (2018). A. Left hemisphere; B. Right hemisphere; C. Left hemisphere inflated; D. Right hemisphere inflated. IFJa – anterior inferior frontal junction; IFJp – posterior inferior frontal junction.

Results from task-based neuroimaging and patient studies provide compelling evidence that the IFJ supports all three domains of cognitive control. Regarding working memory, robust IFJ activation is commonly observed when subjects actively represent and update the rules for a given trial (Brass & Cramon, 2004; Derrfuss et al., 2005). Because these task goals in working memory must be dynamically adjusted, the IFJ is also highly sensitive to demands on task-switching or cognitive flexibility, and is indeed recruited across stimulus switching, response switching, and cognitive set switching tasks (Kim et al., 2011). The IFJ also co-activates with regions of the frontoparietal, salience, and cingulo-opercular networks during fMRI tasks of task-switching in adults (Armbruster et al., 2012). Finally, during inhibitory control, the IFJ exhibits consistent activation across diverse inhibition paradigms—an effect that is notably modulated by concurrent working memory load (Barber et al., 2013; Cieslik et al., 2015; Levy & Wagner, 2011).

Further, evidence from clinical patient populations suggests that IFJ is not just correlated with—but causally necessary for—cognitive control. Positron emission tomography and lesion studies reveal that hypometabolism or damage to the IFJ yields deficits across all three domains of cognitive control, resulting in profound clinical functional impairments including behavioral disinhibition, planning deficits, and impaired problem solving (Aron et al., 2003; Knutson et al., 2015; Schroeter et al., 2012, 2020).

### Circuit architecture of a functional hub for cognitive control

The IFJ demonstrates both anatomical and functional specificity relative to neighboring precentral and frontal sulcal regions. Structurally, it possesses a unique laminar makeup relative to the rest of inferior frontal sulci. IFJ contains a densely packed granular layer II, and large pyramidal cells in deep layer III and upper layer V (Ruland et al., 2022). These features leave IFJ well-suited to optimize large-scale functional network connectivity changes required for the implementation of cognitive control. In other words—IFJ’s laminar structure would enable it to integrate complex, multi-areal inputs via its dense granular layer II, and drive far-reaching shifts in cortico-cortical and cortico-subcortical connectivity via its large pyramidal cells in layers III and V, respectively. This circuit architecture is optimized for the rapid initiation of top-down control.

Functionally, while the IFJ is reliably recruited across inhibitory control, cognitive flexibility, and working memory tasks, its neighboring structures do not show this level of domain-general involvement in cognitive control. At the caudal extent of IFJ, the neighboring PEF region of the IPCS is primarily involved in oculomotor control during saccades, and is functionally connected to the dorsal attention network (Bedini et al., 2023; Bedini & Baldauf, 2021; Derrfuss et al., 2012; Schaefer et al., 2018). At IFJ’s rostral extent, the neighboring inferior frontal sulcus (IFS; IFSp in Glasser et al., 2016) and inferior frontal gyrus (IFG; area 44 in Glasser et al., 2016) regions play a specialized role in the detection of cues to inhibit prepotent motor responses (Cai & Leung, 2011). Right IFG, especially, is thought to represent the core PFC node of a hyperdirect frontostriatal pathway for rapid response inhibition (Aron et al., 2016).

The IFJ’s unique anatomical and functional circuit architecture make it ideally-suited to act as a functional hub between frontoparietal, default, cingulo-opercular, ventral attention/salience, and dorsal attention networks (Asplund et al., 2010; Dixon et al., 2018; Yin et al., 2018). Task-based changes in functional connectivity of the IFJ are observed depending on cognitive demands—with IFJ coupling with ventral attention/salience network during exogenously-cued attention tasks and dorsal attention network during endogenously-oriented attention tasks (Asplund et al., 2010; Dixon et al., 2018). Further, the left IFJ acts as a flexible bridge between frontoparietal and cingulo-opercular (aka, “action mode”; Dosenbach et al., 2025) networks during rule switches, with effective connectivity of these networks to the IFJ predicting individual differences in behavioral assays of cognitive flexibility (Yin et al., 2018). The IFJ has also been found to exert far-reaching modulatory influences on anterior cingulate cortex, parietal regions, and subcortical regions (Baldauf & Desimone, 2014; Hippmann et al., 2021; Sneve et al., 2013). Therefore, the IFJ’s specialized structural and functional connectivity profile make it an ideal hub for the coordination and implementation of cognitive control across inhibitory control, cognitive flexibility, and working memory domains. However, the evidence reviewed so far has come entirely from neuroimaging and clinical studies in adults. Therefore, what remains underexamined is the neurodevelopment of IFJ and its functional role in cognition. Understanding how the functional topography of the IFJ matures across these three domains of cognitive control from childhood through adulthood is a critical open question for both cognitive and developmental neuroscience.

### The neurodevelopment of cognitive control

Theories of neurodevelopment associate decreases in risk-taking behaviors found from adolescence to adulthood with the maturation and refinement of cognitive control processes (Crone & Steinbeis, 2017; Tervo-Clemmens et al., 2023). This appears to be shaped by reliable changes in brain structure and function from childhood through emerging adulthood. This includes changes in white matter myelination and axon density (Lebel et al., 2019; Luo et al., 2025), alongside synaptic pruning and thinning in gray matter (Luna et al., 2015), both of which demonstrate precipitous changes in adolescence. In parallel, functional connectivity analyses reveal a neurodevelopmental transition from local, proximity-based co-activation in childhood to distributed, brain-wide functional modules in adulthood (Engelhardt et al., 2019; Fair et al., 2009). These brain-wide neurobiological changes likely enhance the brain’s neural efficiency, allowing for faster communication and improved cognitive control (Engelhardt et al., 2019; Luo et al., 2025).

At the behavioral level, the three domains of cognitive control appear to exhibit partially distinct developmental trajectories. Performance across inhibitory control, cognitive flexibility, and working memory tasks generally improves with age (Tervo-Clemmens et al., 2023), but the timeline and constancy of these maturational curves is a matter of debate. For example, it has been argued that inhibitory control plateaus in late childhood and early adolescence, while working memory and cognitive flexibility continue to mature into late adolescence / early adulthood (Cowan, 2016; Cragg, 2016; Luna et al., 2004). In contrast, the largest study to date on the maturation of behavioral markers of cognitive control in youth appears to suggest a more uniform, parallel developmental trajectory across domains (Tervo-Clemmens et al., 2023). The ambiguity of developmental findings in the behavioral literature underscores the need for a meta-analytic approach examining the neural substrates of cognitive control across development—to directly test whether cognitive control develops monolithically, or through distinct, domain-specific neurodevelopmental changes.

There is a relative lack of data on the neurodevelopment of IFJ or its role in cognitive control domains in youth. The few existing neurodevelopmental studies focused on IFJ report mixed findings regarding age-related changes in its activation and functional connectivity. Studies have reported both stable activation of IFJ across children and adults during cognitive control tasks, alongside increased connectivity between IFJ and prefrontal and parietal regions—suggesting less up-regulation of task-related activation via IFJ connectivity might be taking place in youth compared with adults during cognitive control (Morton et al., 2009; Schwarze et al., 2023). Given the potential functional utility of the IFJ in cognitive control and network communication, understanding how its flexible recruitment across the different domains changes over development provides a unique insight into the neurobiological basis of the maturation of cognitive control.

### Our approach: A coordinate-based neuroimaging meta-analysis

Here, we conducted a series of coordinate-based neuroimaging meta-analyses to provide further evidence for the domain-general involvement of IFJ in cognitive control in adults, and to characterize the neurodevelopmental trajectory of IFJ involvement in different domains of cognitive control in youth relative to adults. Specifically, we aimed to test the following three *a priori* hypotheses:

***1) IFJ is a domain-general hub for cognitive control in adults.*** The IFJ will be activated across working memory, cognitive flexibility, and inhibitory control tasks in adults.
***2) IFJ flexibly changes its co-activation pattern depending on control demands in adults.*** Different cognitive control domains will demonstrate distinct co-activation patterns of IFJ with other task-positive brain networks including frontoparietal network, cinguloopercular/action mode network, ventral attention/salience network, and the dorsal attention network.
***3) Domain-specific trajectories of IFJ neurodevelopment in cognitive control. IFJ involvement in cognitive control will*** be enhanced in adults relative to youth across inhibitory control, cognitive flexibility, and working memory domains.

These hypotheses will be tested using the Neurosynth Compose tool (J. Kent et al., 2024; J. D. Kent et al., 2026). Hypotheses 1 and 2 will be tested using multilevel kernel density analyses (MKDAs) of existing adult neuroimaging studies in the three cognitive control subdomains. Hypothesis 3 will be evaluated via MKDA Chi^2^ tests to compare activation maps across adult and youth fMRI studies in each of the cognitive control subdomains. We will then discuss the implications of the current meta-analysis for understanding the neurodevelopment of cognitive control, and the functional role of the IFJ in cognitive control more broadly.

## Methods

A series of coordinate-based meta-analyses were conducted to examine the functional maturation of cognitive control by examining fMRI activation in adults during measures of the three domains of cognitive control: working memory, task-switching (i.e., the most commonly used paradigm for probing cognitive flexibility), and inhibitory control. Additionally, three Chi^2^ analyses were run to examine activation differences in youth and adults during measures of the each of the domains. The Neurosynth Compose platform was utilized to curate the search results, extract the coordinates from included articles, as well as specify and run the analyses (J. Kent et al., 2024; J. D. Kent et al., 2026). All analyses were executed by the Neuroimaging Meta-Analysis Research Environment (NiMARE) v0.8.0 (RRID:SCR_017398) Python package (Salo et al., 2022, 2023). The IFJ was defined using the Glasser atlas, which as aforementioned, parcellates the region into the IFJa and IFJp subregions (Glasser et al., 2016).

### Study identification, screening, and inclusion/exclusion

PubMed and Google scholar online databases were searched with terms related to neuroimaging, inhibitory control, switching, and working memory, and well as terms which specified the developmental period (i.e., adult or youth; See Supplemental Materials for search terms). The search results were uploaded to individual projects on the Neurosynth Compose website (J. Kent et al., 2024; J. D. Kent et al., 2026) where each of the study sets were curated according to the Preferred Reporting Items for Systematic reviews and Meta-Analyses (PRISMA) guidelines (See **Figure 2**): 1) duplicates were first identified and removed; 2) article titles and abstracts were then screened to identify relevant peer-reviewed manuscripts; 3) finally, full texts were reviewed to determine eligibility for inclusion.

**Figure 2.**
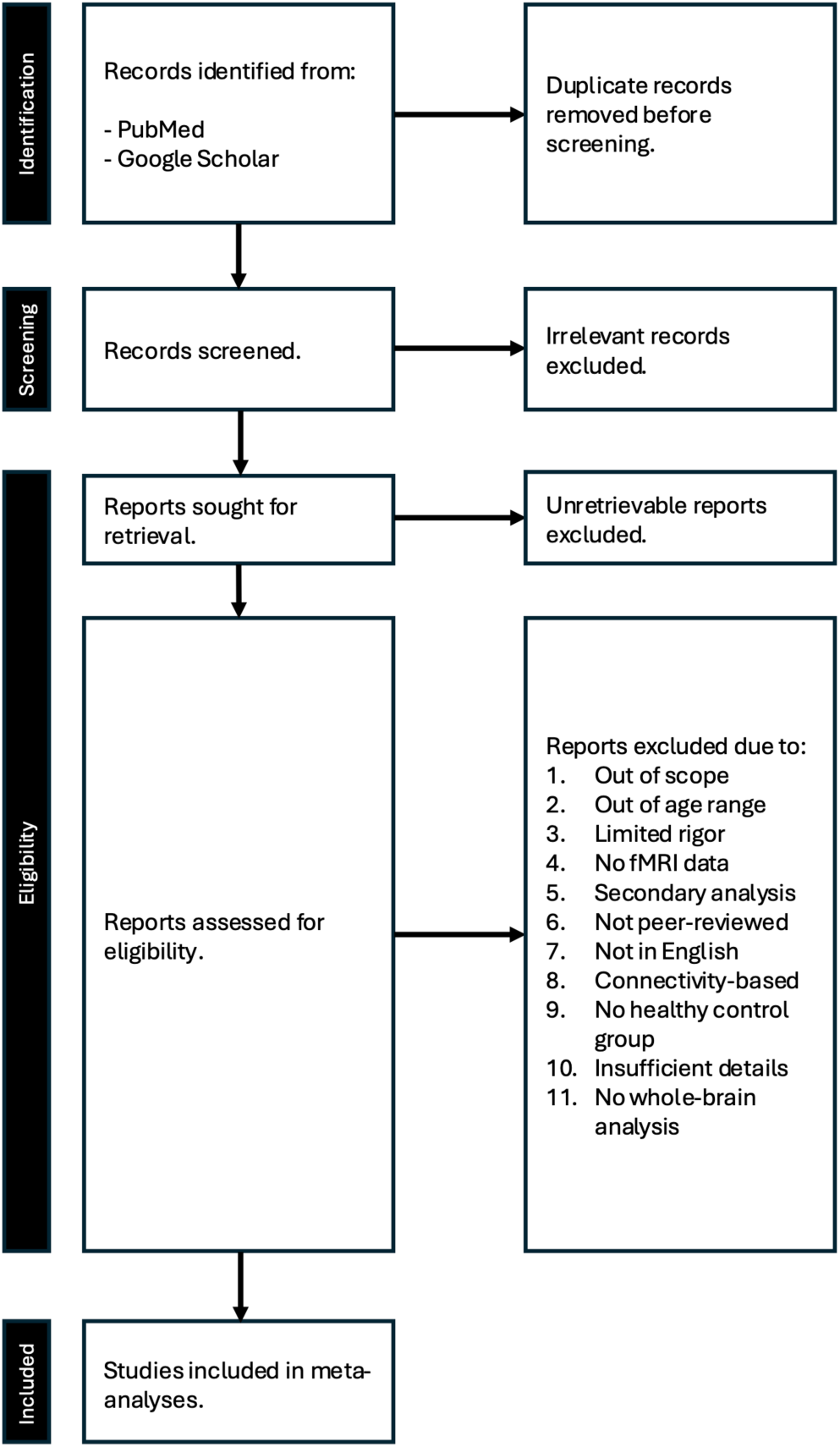
PRISMA diagram of the study curation procedures.

We conducted the original search from any time up to August 2025 and conducted another search to update the results in January 2026. These searches identified a total of 17,093 in children and adolescents (Pubmed: 15,448; Google Scholar: 1,645), and 7,677 studies in adults (Pubmed: 4,820; Google Scholar: 2,857). 565 duplicates were removed from the studies in youth, and 272 duplicate articles were removed from the search results in adults. 6087 studies in youth and 3764 studies in adults were removed after being screened for relevancy, 2 reports were not retrieved for youth, and 5 were not retrieved in adults, leaving 10,439 studies in youth and 3,759 studies in adults to be screened for inclusion criteria (See **Supplemental Figures 1-6** for PRISMA diagram for each meta-analysis). The remaining studies were screened according to the following inclusion criteria:

1. All manuscripts must be peer-reviewed and published in English.
2. All studies must include task-based fMRI, and MNI or Talairach coordinates must be available.
3. The studies must include healthy samples and report those contrasts without the inclusion of other groups.
4. All manuscripts must report the age of the samples included.
5. Whole-brain analyses must be available as ROI-based analyses would inflate the significance for those regions included.

Hence, non-peer-reviewed manuscripts (e.g., abstracts, book chapters, dissertations, theses, etc.) and case-studies were excluded. Studies only reporting on non-healthy samples, and samples with participants who are out of the age ranges (youth: 6-18 years; adults: >18-45 years) were also excluded. Moreover, connectivity-based analyses were excluded. Genotype studies and studies which include participants of a single gender were considered to be of “limited rigor” as they are not representative of the larger population and were therefore excluded from the current analyses as well.

### Cognitive Control Measures

fMRI tasks that measure working memory such as *n*-back tasks, as well as verbal and visual working memory tasks were included in the current meta-analysis. The Sternberg Working

Memory/Recognition task has been argued to be a measure of recognition or short-term memory, as opposed to true working memory (Corbin & Marquer, 2013), so was excluded from the analyses. Several fMRI tasks are considered to measure both cognitive and motor aspects of inhibition; the Go/No-go, Stroop, Stop Signal, Flanker, Anti-saccade, Multi-source interference, and Response-inhibition tasks were included to capture this range of inhibitory control processes. Shifting or switching has been measured with task-switching paradigms, set-shifting, rule-switching, and Wisconsin Card Sorting tasks, all of which were included in the current meta-analyses. See **Supplemental Table 1** for task descriptions.

The current meta-analysis was focused on understanding the neurodevelopment of “cold” cognitive control mechanisms, therefore tasks with affective or motivational manipulations were excluded (e.g. tasks involving emotional faces or reward-, pain-, drug-, and food-related cognitive control cues). We also focused on tasks involving visual cues, as this is the most common sensory modality used in this field (i.e., tasks involving auditory, spatial, and tactile cues were excluded). These distinct affective and multisensory components may elicit activity in distinct networks (Fascher et al., 2026; Mansueto et al., 2025; Sevel et al., 2026; N. Wang et al., 2026; Q. Zhang et al., 2025), and are beyond the scope of the current meta-analysis. There are switching paradigms with other components as well, such as language switching tasks, perspective switching tasks, and voluntary switching tasks which are considered out of the scope of this manuscript as their task designs are optimized to probe distinct processes beyond cognitive flexibility, and therefore they likely recruit distinct and idiosyncratic control networks (Czaczkes et al., 2018; Kousaie & Klein, 2025; Y. Wang et al., 2015).

### Inclusion Summary

According to the criteria, 18 studies in adults and 5 in youth were included in the inhibition analyses, 23 studies in adults and 2 in youth were included in the switching analyses, and 17 studies in adults and 6 in youth were included in the working memory analyses (see **Table 1**). Once the study sets were curated, coordinates for the contrasts of interest were entered into the Neurosynth Compose project, which transforms Talairach coordinates to MNI space. There were a total of 1,285 adult and 628 youth participants, 624 coordinates from the adult studies, and 155 coordinates from the studies in youth.

**Table 1.**
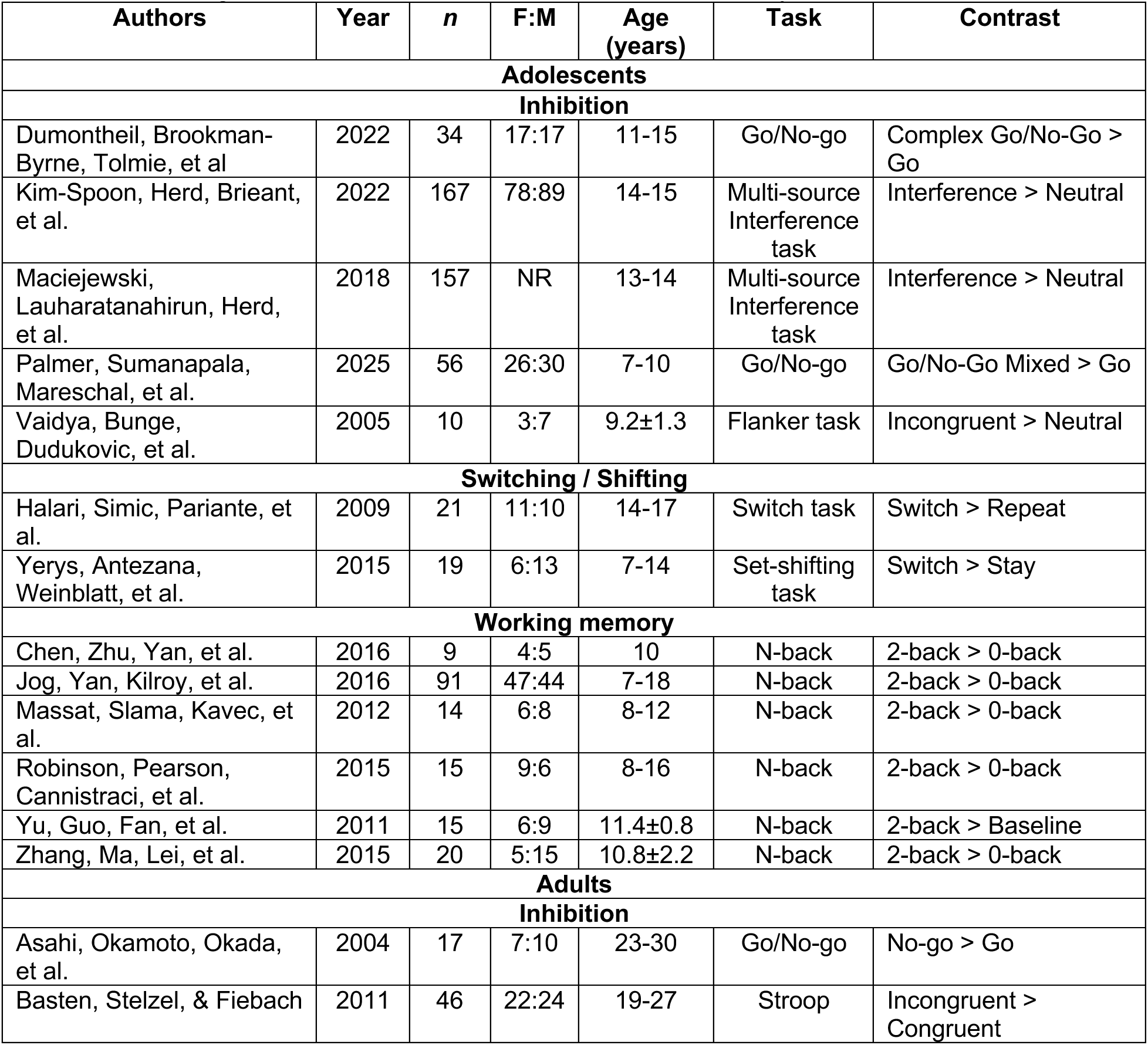
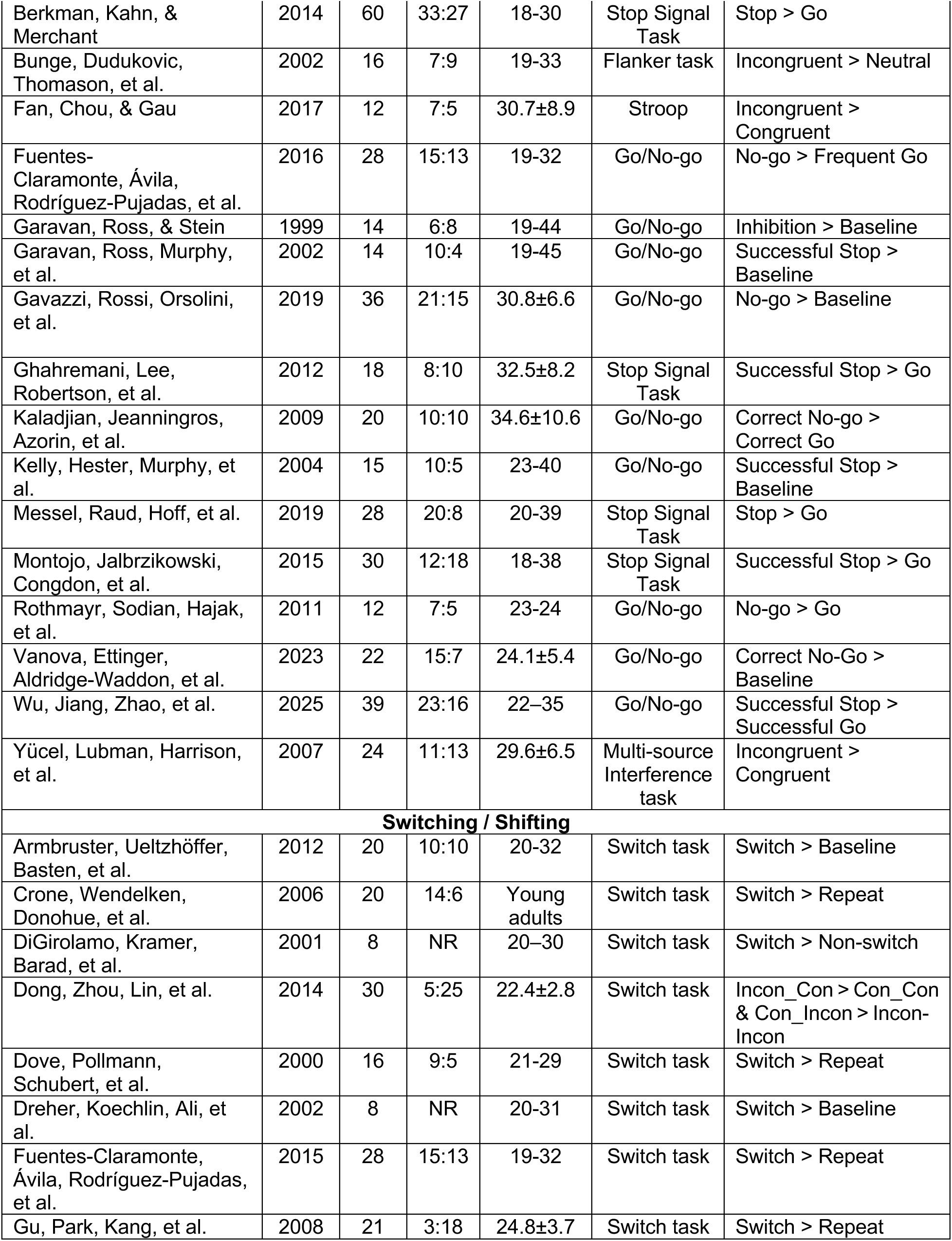
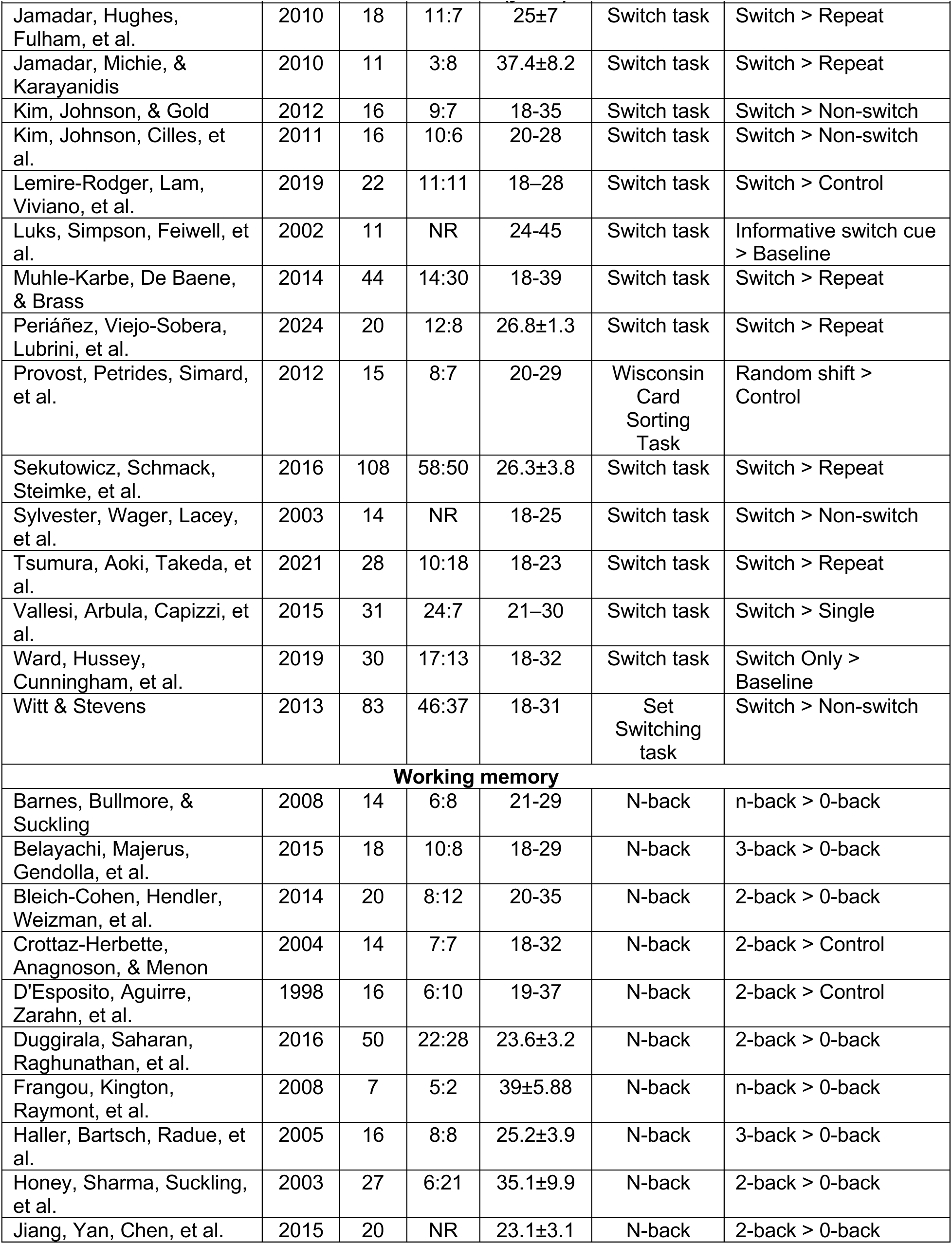
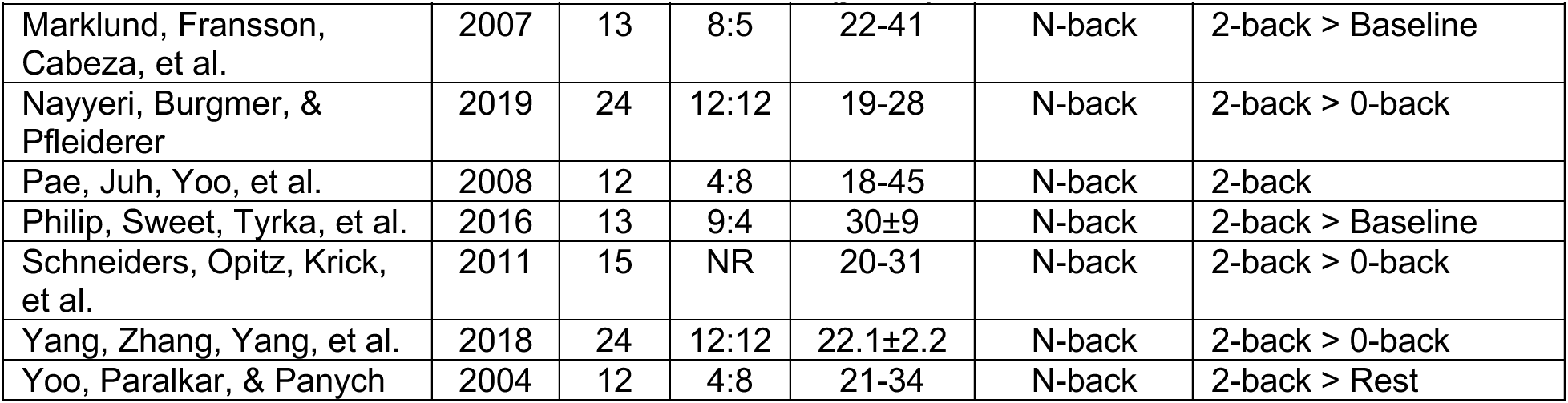
Table listing all studies and contrasts included in the meta-analyses.

### Meta-analyses procedures

Six coordinate-based meta-analyses were conducted using the MKDA approach (Wager et al., 2009) available on the Neurosynth Compose website (J. Kent et al., 2024; J. D. Kent et al., 2026). For both youth and adults, during each of the three domains of cognitive control, peak coordinates of active foci were nested in study contrast maps. These maps were treated as random effects, and used as the unit of analysis (Kober & Wager, 2010; Wager et al., 2009). In this kernel method, each coordinate is convolved with a sphere with a radius of 10.0 and a value of 1; for voxels with overlapping spheres, the maximum value was retained (J. Kent et al., 2024; J. D. Kent et al., 2026). Summary statistics were converted to p-values using an approximate null distribution (J. Kent et al., 2024; J. D. Kent et al., 2026). A Family Wise Error (FWE) correction was utilized – this method uses Monte Carlo simulations to obtain a threshold and establish statistical significance (Kober & Wager, 2010; Wager et al., 2009). In this procedure, null datasets are generated in which dataset coordinates are substituted with coordinates randomly drawn from the meta-analysis mask, and maximum values are retained (J. Kent et al., 2024; J. D. Kent et al., 2026). This procedure was repeated 5000 times to build null distributions of summary statistics, cluster sizes, and cluster masses. Clusters for cluster-level correction were defined using edge-wise connectivity and a voxel-level threshold of p < 0.001 from the uncorrected null distribution (J. Kent et al., 2024; J. D. Kent et al., 2026).

Three MKDA Chi^2^ tests were also conducted using Neurosynth Compose (J. Kent et al., 2024; J. D. Kent et al., 2026). This Chi^2^ method is a test of independence which offers the allowance to test absolute differences in activation between conditions or groups (Kober & Wager, 2010; Wager et al., 2009). It compares observed activation frequencies of comparison indicator maps from individual MKDAs with the null hypothesis of equal expected frequencies across all groups to compare activation in any given voxel (Kober & Wager, 2010; Wager et al., 2009). As with the individual MKDAs, a 10mm spherical kernel was used, for voxels with overlapping spheres, the maximum value was retained. An FWE correction with 5000 iterations of the Monte Carlo procedure was conducted with a voxel-level threshold of p < 0.001.

## Results

Through a series of neuroimaging meta-analyses, we aimed to characterize the activation and co-activation patterns of the IFJ during cognitive control in adults and youth and examine whether there is evidence that this region may act as a functional hub between networks. We conducted six individual MKDAs to investigate activation in adults and youth during inhibitory control, switching, and working memory cognitive control domains, and three MKDA Chi^2^ analyses to examine age related differences during each domain. In adults, convergence of activation was found in the left IFJ during tasks of switching and working memory, as well as in ventral premotor regions, bordering the right IFJ, during inhibition. In youth, convergence of activation was found bilaterally in the IFJ during inhibition, but not during switching or working memory, which when compared with adult activation patterns, suggests ongoing specialization of the region for cognitive control processes during development. The results of each analysis are described in detail below:

### Inhibition

#### MKDA: Adults

The MKDA on inhibition in adults included 18 studies, 241 foci, and 480 participants, and revealed 4 clusters (see **Figure 3A** & **Table 2**). A cluster bordering the right IFJ including the right dlPFC was found, convergence of activation was also found in the pre-SMA (bilateral), and in the right anterior insula. No convergence of activation in or around the left IFJ was found. Given that the dlPFC and pre-SMA are both nodes in the FPN, these results largely replicate previous findings in adults of concurrent recruitment of the FPN and CON (Armbruster et al., 2012), though do not provide support for consistent convergence of activation in the IFJ specifically.

**Figure 3.**
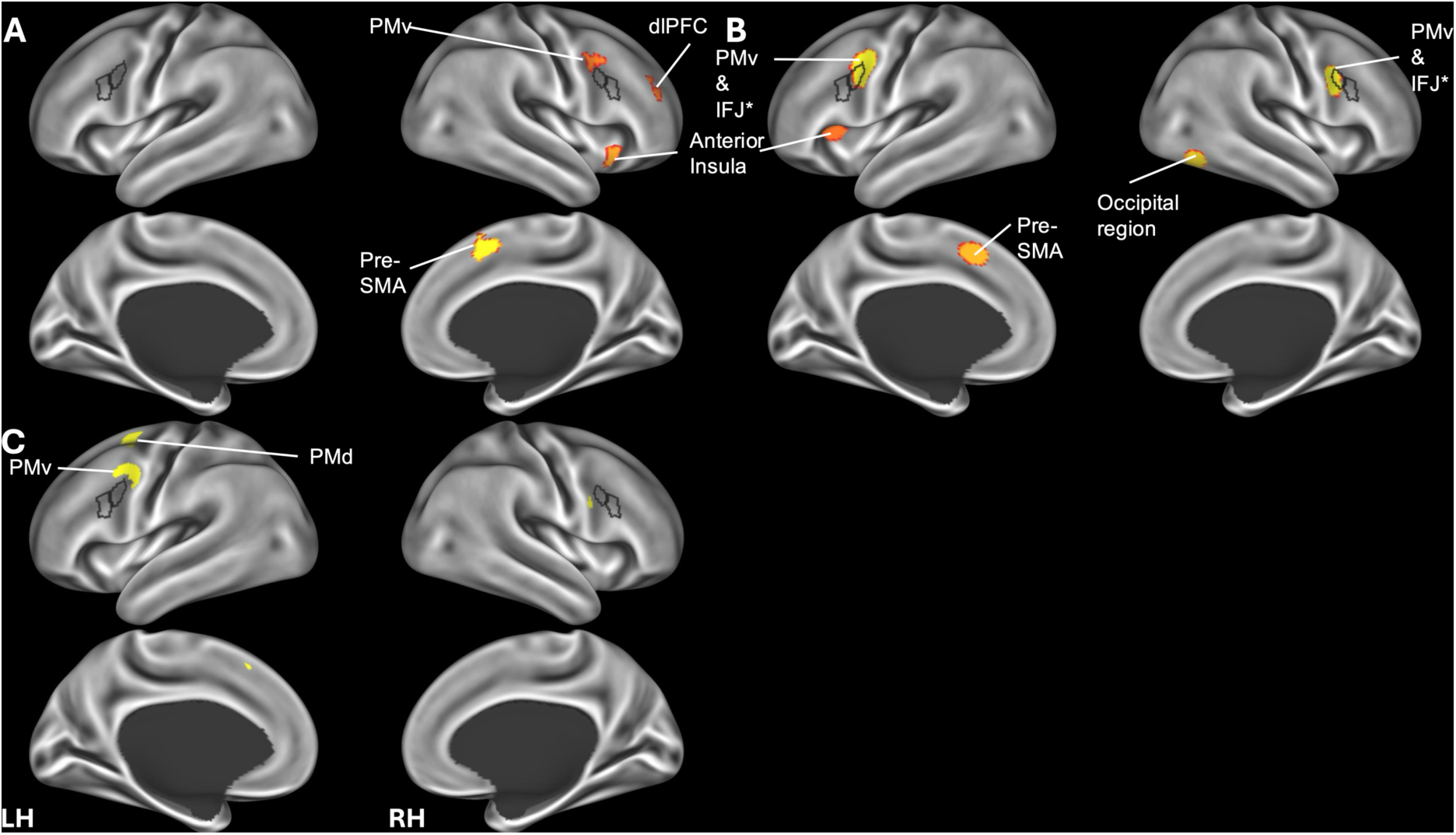
Inhibition. A. Adult MKDA results; B. Youth MKDA results; C. MKDA Chi^2^ results. Voxel-level threshold = 1.65. Acronyms: dlPFC, dorsolateral prefrontal cortex; PMd, dorsal premotor cortex; PMv, ventral premotor cortex; IFJ, inferior frontal junction; SMA, supplementary motor area.

**Table 2.** Results of the individual MKDAs.

| Region | Glasser Area(s) | Hemisphere | X | Y | Z | Peak Z Stat | Cluster Size |
| --- | --- | --- | --- | --- | --- | --- | --- |
| *Adult - Inhibition* |  |  |  |  |  |  |  |
| Pre-Supplemental Motor Area | SCEF, 8BM, SFL, p32pr | Right | 4 | 12 | 50 | 3.54 | 3064 |
| Anterior Insula | AVI, AAIC, MI | Right | 34 | 18 | -6 | 2.85 | 1712 |
| Ventral Premotor Cortex | PEF, 8c | Right | 42 | 6 | 36 | 2.62 | 1040 |
| Dorsolateral Prefrontal Cortex | 46 | Right | 36 | 38 | 24 | 2.59 | 1008 |
| *Youth – Inhibition* |  |  |  |  |  |  |  |
| Inferior Frontal Junction & Ventral Premotor Cortex | IFJp, PEF, 8c, 6r | Left | -44 | 2 | 34 | 3.54 | 3064 |
| Pre-Supplementary Motor Area | SCEF, p32pr, 8BM | Left | -4 | 14 | 48 | 2.91 | 2728 |
| Inferior Frontal Junction & Ventral Premotor Cortex | IFJp, PEF, 6r | Right | 46 | 6 | 28 | 2.85 | 2104 |
| Occipital Lobe | PH, FFC | Right | 42 | -62 | -10 | 2.64 | 2304 |
| Anterior Insula | AVI, FOP4-5 | Left | -30 | 20 | 8 | 2.55 | 2120 |
| *Adult – Switching* |  |  |  |  |  |  |  |
| Dorsolateral Prefrontal Cortex, Inferior Frontal Cortex*, & Ventral Premotor Cortex | IFJa, IFJp, PEF, FEF, 6a, 6r, 6-8, 8c, 55b, p9-46v, p47r, 46, IFSa, IFSp | Left | -40 | 14 | 34 | 3.54 | 16624 |
| Superior Parietal Region | AIP, IP2, 7PC, LiPd, LiPv, MIP, PFm | Left | -36 | -52 | 48 | 3.54 | 7640 |
| Pre-Supplementary Motor Area | SCEF, p32pr, 8BM, a32pr<sup>‡</sup>, SFL<sup>‡</sup> | Bilateral | -2 | 16 | 48 | 3.54 | 6632 |
| Medial Parietal Region | 7Pm, POS2, 7PL | Right | 14 | -68 | 50 | 2.64 | 2400 |
| Anterior Insula Region | AVI, FOP4 | Right | 32 | 22 | 0 | 2.03 | 1784 |
| *Youth – Switching* |  |  |  |  |  |  |  |
| No significant clusters |  |  |  |  |  |  |  |
| *Adult – Working memory* |  |  |  |  |  |  |  |
| Inferior Frontal Junction & Ventral Premotor Cortex | IFJp, 6r, 6v, FEF, PEF, 55b, 8c | Left | -46 | 4 | 36 | 3.54 | 3480 |
| Intraparietal Region | AIP, LiPd, PFm, IP1-2, MIP | Left | -38 | -52 | 46 | 3.54 | 7504 |
| Anterior Insula | AIV, FOP4-5, MI | Left | -30 | 22 | 0 | 3.54 | 3768 |
| Pre-Supplementary Motor Area | SCEF, SFL<sup>‡</sup>, p32pr<sup>‡</sup>, 8BM | Bilateral | 0 | 14 | 48 | 3.54 | 6104 |
| Dorsal Premotor Cortex | 6a, i6-8 | Right | 28 | 6 | 58 | 3.54 | 3824 |
| Anterior Insula | AIV, FOP4-5, MI | Right | 34 | 22 | 2 | 3.54 | 3416 |
| Intraparietal Region | AIP, IP1-2, PFm, LiPd | Right | 40 | -46 | 44 | 3.54 | 5856 |
| Dorsolateral Prefrontal Cortex | 46, p9-46v | Right | 44 | 40 | 20 | 1.67 | 1696 |
| *Youth – Working memory* |  |  |  |  |  |  |  |
| Pre-Supplementary Motor Area | SCEF<sup>‡</sup>, p32pr<sup>‡</sup>, a32pr, 8BM | Bilateral | 0 | 22 | 44 | 3.54 | 3200 |
| Anterior Insula | AVI | Right | 32 | 22 | -4 | 3.24 | 2808 |
| Intraparietal Region | PFM, IP1-2 | Right | 44 | -50 | 46 | 3.16 | 2144 |
| Caudate | - | Left | -16 | -2 | 16 | 2.30 | 1904 |

Coordinates reported in MNI space. \* Indicates cluster includes the Inferior Frontal Junction. <sup>‡</sup> Indicates Glasser area is in the left hemisphere only.

#### MKDA: Youth

The MKDA on inhibition in youth included 5 studies, 79 foci, and 424 participants, and 5 clusters of activation convergence were found. Convergence of activation was found bilaterally in the ventral premotor cortex, IFJp, and a portion of dlPFC (see **Figure 3B** & **Table 2**), co-activation was found with another node in the FPN, the pre-SMA. Convergence was also found in a node of the CON, the left anterior insula, as well as in the right occipital region. These findings indicate the FPN activates along with the DAN and CON during fMRI tasks of inhibitory control in youth, which is largely a replication of what has been reported in the literature, however previous studies have attributed FPN activation to regions other than the IFJ, such as the dlPFC, which also displayed convergence of activation in the current analysis, or the medial frontal gyrus, which did not display convergence of activation in the current analysis (Engelhardt et al., 2019).

#### MKDA Chi^2^: Age differences

All included inhibition studies used in the individual MKDAs in youth and adults were used in the MKDA Chi^2^ analyses. Two clusters reached significance (see **Figure 3C** & **Table 3**), one in the left premotor cortex adjacent to left posterior IFJ, which includes the PEF and a portion of the dlPFC (area 8c). Another cluster covers the left parietal cortex and includes the premotor region. The results suggest that the IFJ undergoes hemispheric specialization during development whereby bilateral activity in the IFJp during inhibitory control decreases and right-lateralized recruitment of frontal regions increases by adulthood.

**Table 3.** Results from the MKDA Chi^2^ tests.

| Region | Glasser Area(s) | Hemisphere | X | Y | Z | Peak Stat | Cluster Size |
| --- | --- | --- | --- | --- | --- | --- | --- |
| *Youth vs Adults – Inhibition* |  |  |  |  |  |  |  |
| Ventral Premotor Cortex | PEF, 8c, 55b | Left | -44 | 0 | 38 | 1.99 | 872 |
| Dorsal Premotor Cortex | 6a, FEF | Left | -26 | 0 | 56 | 1.7 | 672 |
| *Youth vs Adults – Switching* |  |  |  |  |  |  |  |
| Results invalid |  |  |  |  |  |  |  |
| *Youth vs Adults – Working Memory* |  |  |  |  |  |  |  |
| No significant clusters |  |  |  |  |  |  |  |

Coordinates reported in MNI space.

### Switching

#### MKDA: Adults

The MKDA on switching in adults included 23 studies, 389 foci, and 619 participants; 5 clusters were found (see **Figure 4A** & **Table 2**). Convergence of activation was found in several left-lateralized prefrontal regions including IFJp and IFJa, along with other regions of the FPN, such as the pre-SMA, and the left and right parietal cortices, which include the intraparietal sulci, and regions considered to be part of the default network, such as the angular gyrus. IFJ co-activation with the CON is evidenced by the convergence of activation in the right anterior insula. No convergence of activation was found in or around the right IFJ. The current findings show a distributed recruitment of networks, with the FPN co-activating with the cingulo-opercular, default, and dorsal attention networks, replicating previous findings in adults (Armbruster et al., 2012).

**Figure 4.**
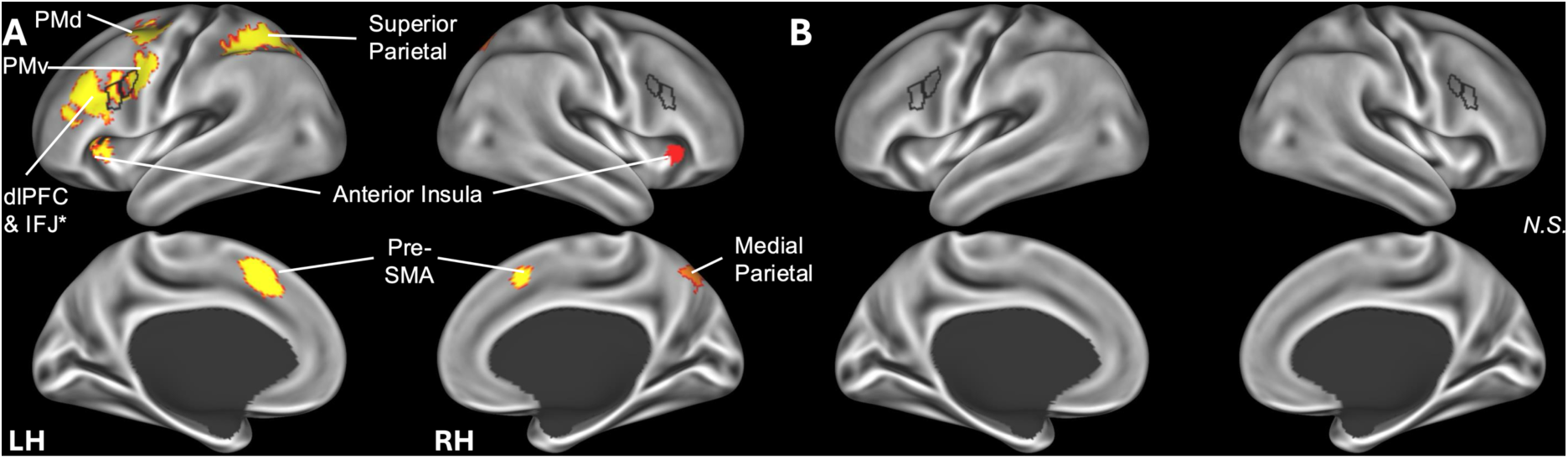
Switching. A. Adult MKDA results; B. Youth MKDA results. Voxel-level threshold = 1.65. Acronyms: dlPFC, dorsolateral prefrontal cortex; PMd, dorsal premotor cortex; PMv, ventral premotor cortex; IFJ, inferior frontal junction; SMA, supplementary motor area.

#### MKDA: Youth

The MKDA on switching in youth included 2 studies, 22 foci, and 40 participants. No significant clusters of convergence were found (see **Figure 4B**). As there were only 2 studies that met the inclusion criteria for the analysis, the current results are presented as exploratory and indicate more studies in youth are needed.

#### MKDA Chi^2^: Age differences

Given the small number of switching studies in youth the results of the MKDA Chi^2^ analysis are considered invalid.

### Working memory

#### MKDA: Adults

The MKDA on working memory in adults included 17 studies, 216 foci, and 315 participants, with 8 clusters found (see **Figure 5A** & **Table 2**). Convergence of activation across the FPN was revealed, specifically in a cluster in left prefrontal and premotor cortices that included left IFJp, another cluster bilaterally across the pre-SMA, as well as in the right dlPFC. Co-activation was found with bilateral clusters containing the inferior parietal cortices, containing the intraparietal sulcus, another node in the FPN and the DAN, as well as the angular gyrus, part of the default network. Convergence was also found bilaterally in the anterior insula, a node in the CON, as well as the Premotor area. No convergence of activation was found in or around the right IFJ for adults during working memory tasks. These results also show distributed network recruitment across the frontoparietal, default, dorsal attention, and cingulo-opercular networks, supporting previous findings (Friedman & Robbins, 2022).

**Figure 5.**
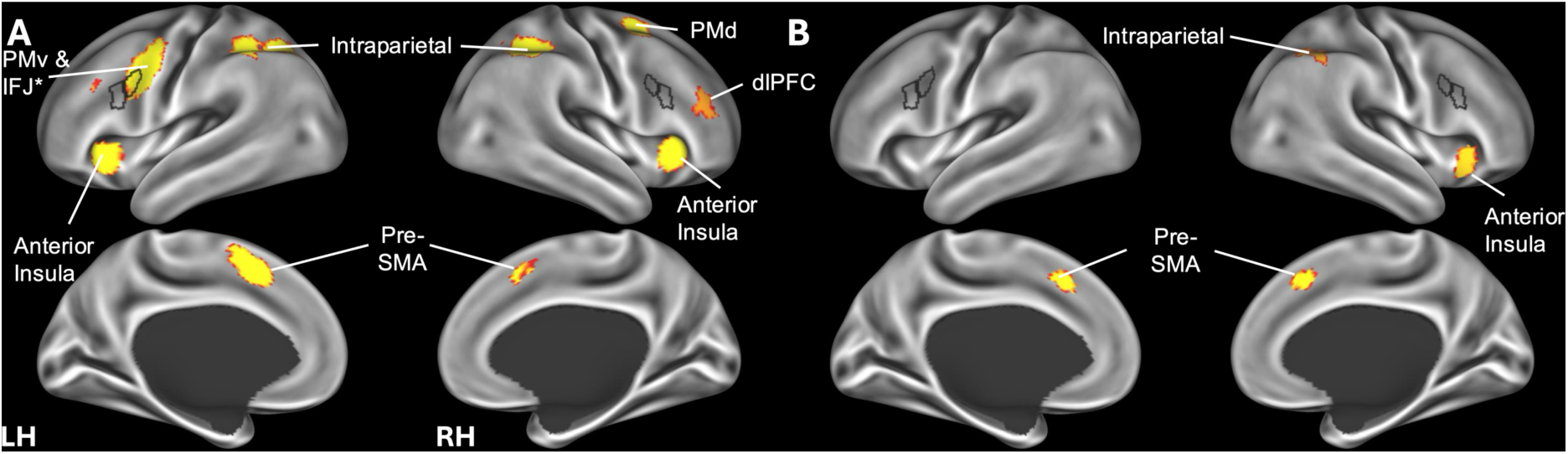
Working memory. A. Adult MKDA results; B. Youth MKDA results. Voxel-level threshold = 1.65. Acronyms: dlPFC, dorsolateral prefrontal cortex; PMd, dorsal premotor cortex; PMv, ventral premotor cortex; IFJ, inferior frontal junction; SMA, supplementary motor area.

#### MKDA: Youth

The MKDA on working memory in youth included 6 studies, 54 foci, and 164 participants. 4 clusters were found including one cluster containing the pre-SMA and extending to the anterior cingulate, both are nodes in the FPN. Convergence of activation was also found in the right inferior parietal cortex, which harbors the intraparietal sulcus, considered to be part of the FPN and the DAN. Additionally, convergence of activation in the right anterior insula, a node in the CON, was revealed. The left caudate, a part of the striatum, also showed convergence during working memory. No convergence in activation in or around the IFJ was found in either hemisphere. (See **Figure 5B** & **Table 2**). These findings show that nodes in the FPN activate along with nodes in the cingulo-opercular and dorsal attention networks during working memory tasks in youth, consistent with previous findings by Engelhardt and colleagues (2019).

#### MKDA Chi^2^: Age differences

All included working memory studies used in the individual MKDAs in adults and in youth were used in the MKDA Chi^2^ analyses. No clusters reached significance, indicating no significant differences in IFJ activation between adults and youth during working memory tasks. This could be indicative of a differential developmental trajectory for working memory processes compared to inhibitory control.

## Discussion

### Meta-Analytic IFJ Signatures of Cognitive Control in Adults

The current MKDAs reliably implicated the IFJ in adult cognitive control. However, closer examination of these results revealed distinct, domain-specific signatures of IFJ recruitment, pointing to both hemispheric and sub-areal specialization. While inhibitory control primarily engaged the right IFJ and adjacent right premotor and dorsolateral prefrontal regions—aligning with established models of right frontal cortex involvement in inhibitory control (Aron et al., 2004, 2007; Rubia et al., 2003)—switching and working memory tasks reliably localized to the left IFJ. Within this left-hemisphere cluster, sub-regions displayed more nuanced activation profiles. For instance, the left IFJp (Glasser et al., 2016), which sits at the functional intersection of the FPN and DAN (Schaefer et al., 2018), showed convergent activation for both switching and working memory. By contrast, recruitment of the left IFJa (Glasser et al., 2016)—a region belonging exclusively to the FPN—converged during switching but not working memory. Across all domains, the IFJ remained robustly coupled with core task-positive networks (FPN, DAN, and CON) and showed striking consistency with the anterior insula, whereas evidence for co-activation with the ventral attention network (VAN) was sparse.

Beyond regional convergence at the IFJ, our results identified clear overlap in the co-activated networks for working memory and task switching, whereas inhibitory control demonstrated a more unique co-activation profile. In inhibitory control, the right-lateralized premotor regions co-activated with right dorsolateral prefrontal cortex (dlPFC; area 9-46), as well as bilateral anterior insula and pre-SMA. Switching engaged a broader, left-lateralized network comprising IFJa, IFJp, dlPFC (area 8), intraparietal sulcus, premotor and frontal eye fields, bilateral anterior insula, and pre-SMA. Similarly, working memory recruited the left IFJp, bilateral dlPFC (area 9-46), premotor and frontal eye fields, intraparietal sulcus, pre-SMA, and anterior insula.

To be clear, the regions most reliably activated across all domains—providing evidence for a neural “unity” of cognitive control—were the dlPFC, pre-SMA, and anterior insula. The dlPFC has long been linked to cognitive control. Seminal discoveries in nonhuman primates established its fundamental role in maintaining task-relevant information and encoding abstract goals (Goldman-Rakic, 1995; Miller & Cohen, 2001). Contemporary human neuroscience has built upon this foundation, suggesting the dlPFC represents a key node of a hierarchical control network involved in managing abstract goals and temporal contingencies (Badre & Desrochers, 2019; Badre & Nee, 2018). Likewise, the pre-SMA plays a key role in planning goal-directed actions, explaining its consistent engagement across tasks (Nachev et al., 2007). Finally, while the functional network membership of the anterior insula remains controversial, our MKDA specifically implicated the anteroventral insula (AVI; Glasser et al., 2016) across all domains. Though various parcellations have alternately associated the AVI with the FPN (e.g., Yeo & Schaefer; Ji et al., 2019) or the CON/“action mode network” (Dosenbach et al., 2025; Gordon et al., 2016), this discordance in resting-state connectivity, juxtaposed against the unwavering activation observed in our analyses, identifies the AVI as a domain-general integrative hub. These results suggest that the AVI’s functional profile is best captured not by intrinsic functional connectivity, but by task-based fMRI.

### Neurodevelopment of Cognitive Control in Youth

Due to insufficient studies in the youth switching MKDA, our developmental comparison analyses were restricted to the working memory and inhibitory control domains. Here, we observed evidence for a unique, domain-specific pattern of differences between youth and adults between these two domains. For working memory, the youth MKDA demonstrated meta-analytic activation of right AVI, intraparietal sulcus, bilateral pre-SMA, and dorsomedial striatum (caudate). The youth map did not reveal evidence for reliable engagement of IFJ or dlPFC recruitment during working memory tasks, but notably, none of these differences yielded a significant difference between youth and adults in our MKDA Chi^2^ analyses. Overall, the recruitment and specialization of FPN nodes underlying cognitive control across development—i.e., the involvement of IFJ and dlPFC in working memory in adults but not youth—is in line with existing findings in the neurodevelopmental literature (e.g. Engelhardt et al., 2019; Fair et al., 2009).

Striking age-related differences were observed in the inhibitory control domain. Specifically, youth demonstrated bilateral activation in the ventral premotor cortex (region 6a), posterior dlPFC (region 8c), and IFJp, and no meta-analytic activation of anterior dlPFC (area 9-46) during inhibitory control was observed. The MKDA Chi^2^ analysis for inhibitory control revealed significant differences between youth and adults in left-hemispheric inferior frontal activation during inhibition, alongside differential activation in left premotor and parietal regions. These age-related differences support the framework that the IFJ comes “online” for distinct executive processes at different developmental stages. The bilateral IFJp activation seen in youth during inhibition suggests this subregion matures early to support inhibitory demands (Cragg, 2016; Crone & Steinbeis, 2017), before transitioning to the specialized, right-lateralized frontal nodes for inhibitory control commonly observed in adulthood.

Our finding that IFJ’s functional network organization becomes more specialized across development represents a potential neural bases for the behavioral maturation of cognitive control across adolescence. Specifically, a delayed specialization of IFJ may explain why two-factor models of cognitive control, which collapse flexibility unto the working memory and inhibitory control constructs (Karr et al., 2018), has recently been shown to fit pediatric cohorts better than the canonical three-factor model (Miyake et al., 2000). This functional transition raises critical questions about the IFJ’s underlying structural maturation. It is likely that precipitous adolescent changes in white matter myelination and synaptic pruning uniquely enable the IFJ’s dense granular layer II and large pyramidal cells to efficiently integrate multi-areal inputs (Lebel et al., 2019; Luo et al., 2025; Ruland et al., 2022). This structural refinement may then bring the region functionally ‘online’ to support the distributed network-level computations required to enable adult-level performance on cognitive flexibility tasks, in particular.

### Methodological Strengths and Limitations

A primary strength of this investigation is the deployment of the MKDA approach. Compared to traditional Activation Likelihood Estimation (ALE) or standard KDA, MKDA lowers false-positive rates and enhances generalizability by treating study contrast maps as random effects, thus respecting the multilevel nature of the meta-analytic data (Kober & Wager, 2010). Furthermore, MKDA weights outputs by sample size, amplifying the statistical impact of well-powered studies. Some limitations must also be acknowledged. Specifically, the developmental analyses were constrained by a paucity of literature on task-switching or cognitive flexibility that passed our inclusion/exclusion criteria, yielding only two eligible studies and resulting in null MKDA findings for this domain. Expanding the primary literature on cognitive flexibility in youth is critical for future meta-analytic efforts.

### Conclusions and Future Directions

The current meta-analysis re-affirms the IFJ as a core node of the FPN whose subregions exhibit domain-dependent co-activation profiles during cognitive control. A transition of IFJ from a bilateral, distributed activation in youth to lateralized, specialized networks for cognitive control in adulthood reflects the ongoing maturational trajectory of these functions across adolescence—nicely complementing the canonical trajectory of behavioral improvements in cognitive control across this age period (e.g. Tervo-Clemmens et al., 2023). Future investigations must examine the development of IFJ longitudinally to pinpoint critical periods of neuroplasticity and vulnerability. Given the structural variations observed in inferior frontal regions and their association with conditions like ADHD (Derrfuss et al., 2009; Li et al., 2021; Ruland et al., 2022), structure-function coupling analyses are also warranted in this developmental period. Similarly, examining deactivation and functional connectivity (Huang et al., 2024)—particularly considering the high activation variability in the IFJ—will clarify covariation dynamics previously masked by basic co-activation analyses. Ultimately, detailing the structural and functional phenotypes of the IFJ may establish it as a viable biomarker for executive dysfunction, a target for cognitive training, or a site for therapeutic neurostimulation in future studies.

## Acknowledgements

This work was supported by National Institutes of Health through the National Institute on Alcohol Abuse and Alcoholism award number R01AA030283, and the National Science Foundation award number #2237795. We would like the thank Drs. James Cavanagh and Cecilia Hinojosa for helpful comments on an earlier version of this manuscript, as well as Drs. James Kent and Alejandro de la Vega for essential support in using Neurosynth-Compose.

