## Supplementary Materials for "Maturation of Cognitive Control in the Inferior Frontal Junction: A Combined Systematic Review and Coordinate-Based fMRI Meta-Analysis"

*Search terms*

("functional magnetic resonance imaging" OR "fMRI" OR "neuroimaging" AND "adult" AND "working memory"), ("functional magnetic resonance imaging" OR "fMRI" OR "neuroimaging" AND "adult" AND "switching"), ("functional magnetic resonance imaging" OR "fMRI" OR "neuroimaging" AND "adult" AND "inhibitory control"), ("functional magnetic resonance imaging" OR "fMRI" OR "neuroimaging" AND "child*" OR “adolescen*” AND "working memory"), ("functional magnetic resonance imaging" OR "fMRI" OR "neuroimaging" AND "child*" OR “adolescen*” AND "switching"), ("functional magnetic resonance imaging" OR "fMRI" OR "neuroimaging" AND "child*" OR “adolescen*” AND "inhibitory control")

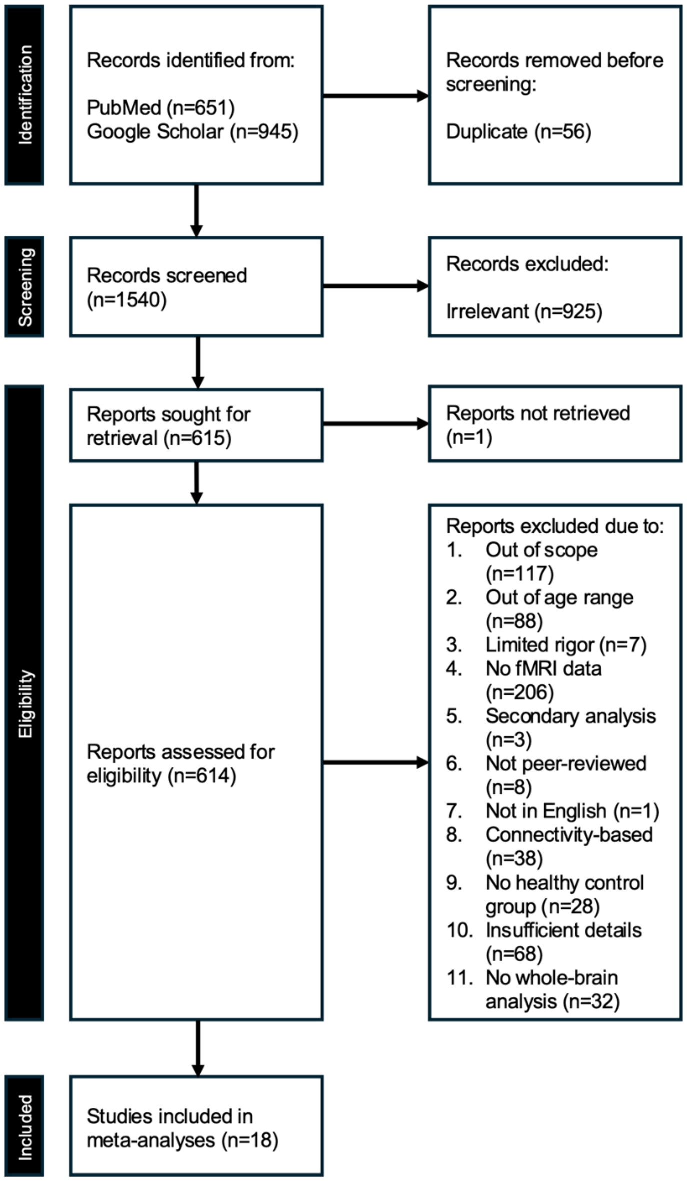


**Supplementary Figure 1.** PRISMA diagram for adults during inhibition

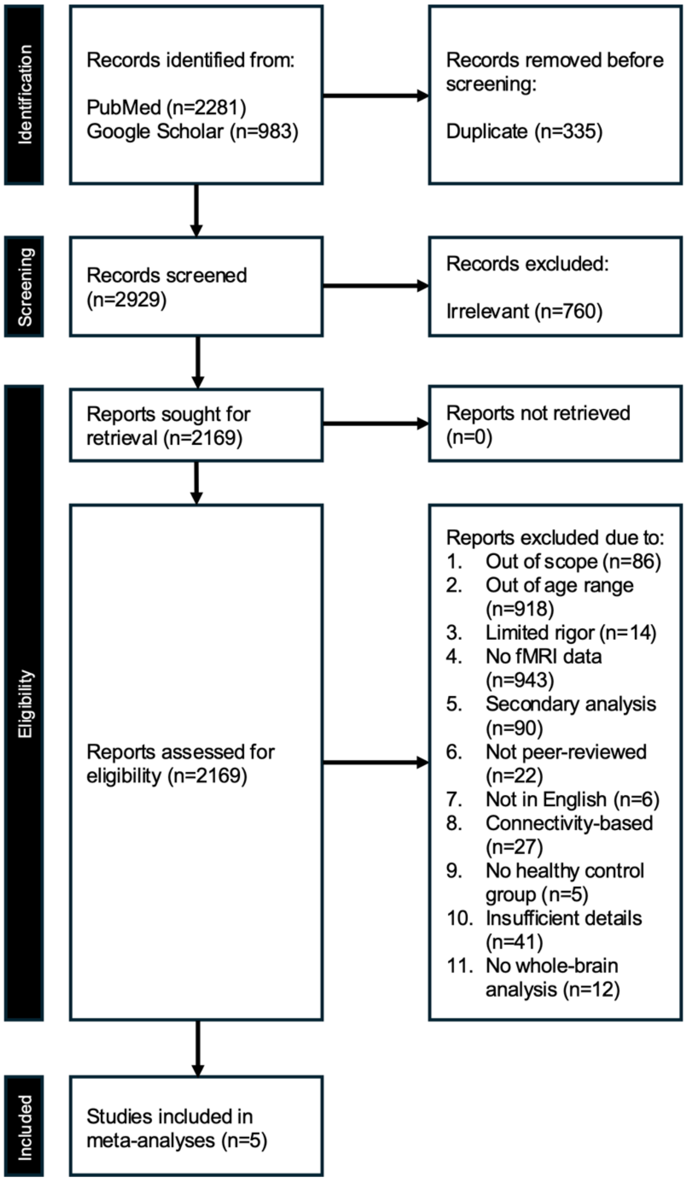


**Supplementary Figure 2.** PRISMA diagram for youth during inhibition

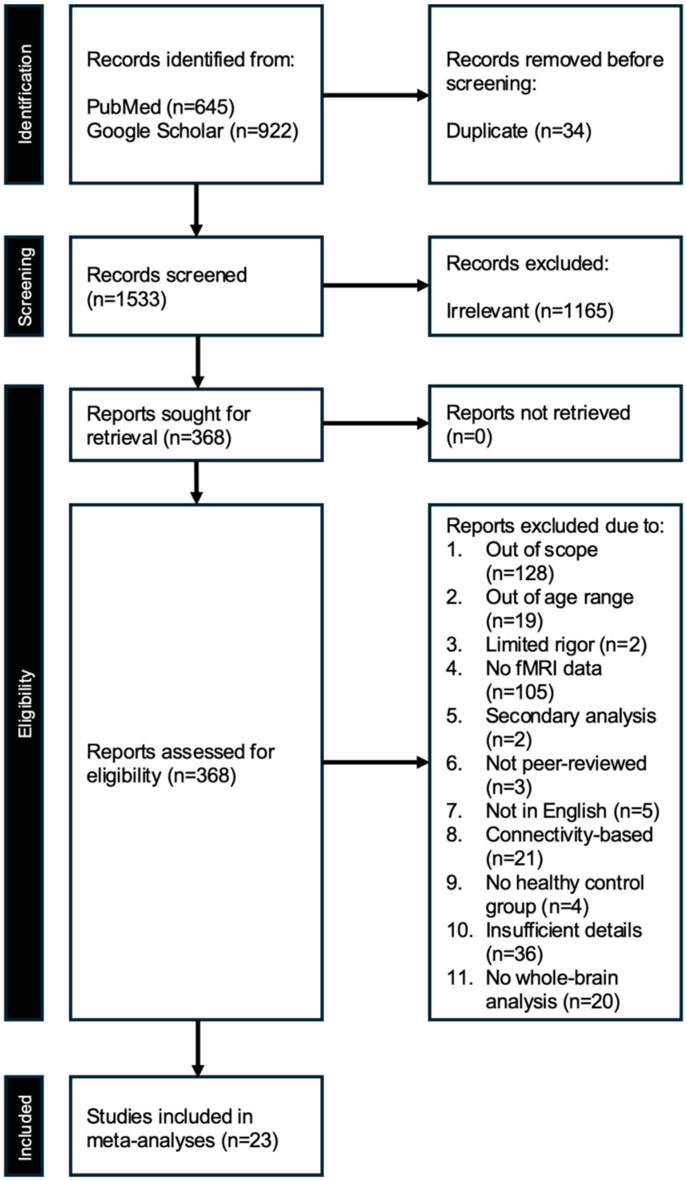


**Supplementary Figure 3.** PRISMA diagram for adults during switching

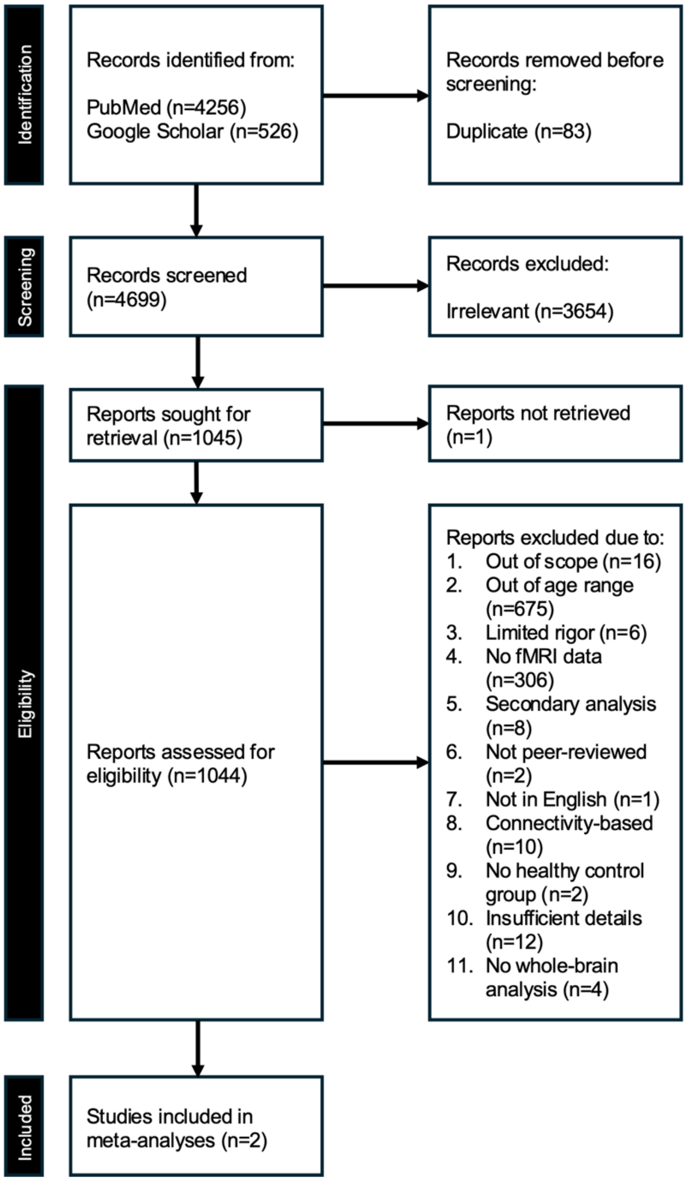


**Supplementary Figure 4.** PRISMA diagram for youth during switching

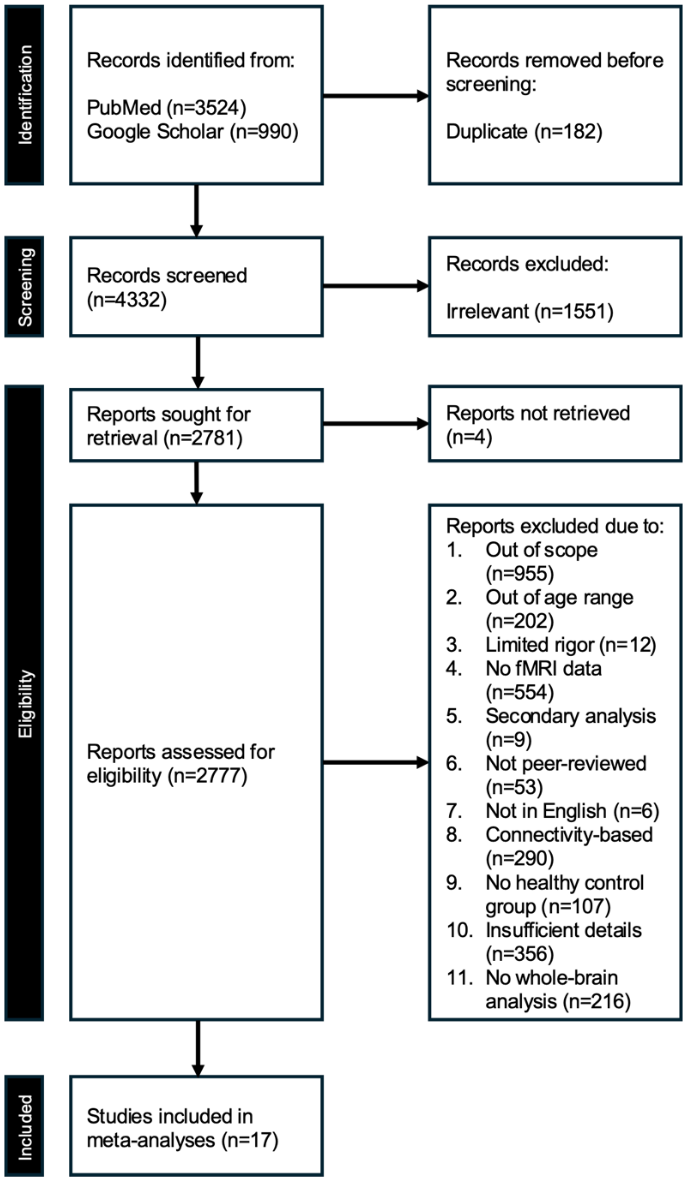


**Supplementary Figure 5.** PRISMA diagram for adults during working memory

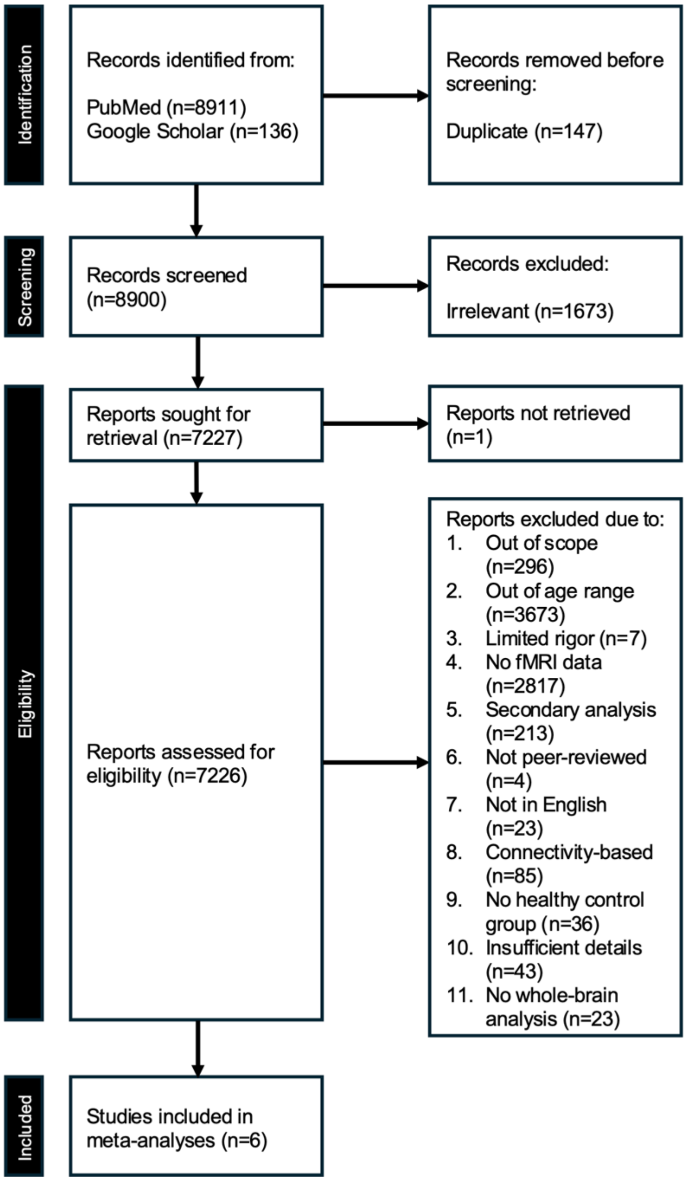


**Supplementary Figure 6.** PRISMA diagram for youth during working memory

**Supplementary Table 1.** Cognitive control task descriptions.

| Domain | Task | Description |
|---|---|---|
| Inhibitory control | Anti-saccade | Participants are asked to fixate on a central stimulus and refrain from moving eyes toward a target stimulus presented in a peripheral location (Magnusdottir et al., 2019). |
| | Flanker | Participants are asked to respond to a central target with distractors on either side (Stins et al., 2007). |
| | Go/No-Go | Participants are asked to respond to a particular cue while withholding their response when presented with a different cue (Raud et al., 2020). |
| | Multi-Source Interference | Participants are asked to respond to a particular cue in the presence of distractor stimuli (Bush et al., 2003). |
| | Stroop | Participants are asked to respond to a specific cue under congruent and incongruent conditions (Parris et al., 2019). |
| | Stop Signal | Participants are asked to respond to the first cue while withholding their response when presented with a stop stimulus (Raud et al., 2020). |
| Switching | Task-switching / Set-shifting | Participants are trained to perform a task based on the specific stimulus presented, when the stimulus changes, they must perform a different task (Kim et al., 2011; Sekutowicz et al., 2016). |
| | Wisconsin Card Sorting | Participants are asked to categorize cards based on a certain feature (Kopp et al., 2021). |
| Working memory | *n*-back | Participants are asked to respond to the stimulus that was presented *n* trials prior (Huang et al., 2025). |
